# Genomics analysis of *Drosophila sechellia* response to *Morinda citrifolia* fruit diet

**DOI:** 10.1101/2021.06.21.449329

**Authors:** Z.A. Drum, S.M. Lanno, S.M. Gregory, Shimshak, W. Barr, A. Gatesman, M. Schadt, J. Sanford, A. Arkin, B. Assignon, S. Colorado, C. Dalgarno, T. Devanny, T. Ghandour, R. Griffin, M. Hogan, E. Horowitz, E. McGhie, J. Multer, H. O’Halloran, K. Ofori-Darko, D. Pokushalov, N. Richards, K. Sagarin, N. Taylor, A. Thielking, P. Towle, J. D. Coolon

**Author notes:** Corresponding Author: Joseph D. Coolon, Wesleyan University, 123 Hall-Atwater, Middletown, CT 06459.

## Abstract

*Drosophila sechellia* is an island endemic host specialist that has evolved to consume the toxic fruit of *Morinda citrifolia*, also known as noni fruit. Recent studies by our group and others have examined genome-wide gene expression responses of fruit flies to individual highly abundant compounds found in noni responsible for the fruit’s unique chemistry and toxicity. In order to relate these reductionist experiments to the gene expression responses to feeding on noni fruit itself, we fed rotten noni fruit to adult female *D. sechellia* and performed RNA-sequencing. Combining the reductionist and more wholistic approaches, we have identified candidate genes that may contribute to each individual compound and those that play a more general role in response to the fruit as a whole. Using the compound specific and general responses, we used transcription factor prediction analyses to identify the regulatory networks and specific regulators involved in the responses to each compound and the fruit itself. The identified genes and regulators represent the possible genetic mechanisms and biochemical pathways that contribute to toxin resistance and noni specialization in *D. sechellia*.

## Introduction

Insects have intimate relationships with plants, ranging from pollination to parasitism, and mimicry to mutualism. One of the most common of these interactions is insect-host plant specialization. A well-studied example of this is the Seychelles Islands endemic fruit fly specialist *Drosophila sechellia* that feeds almost exclusively on the ripe fruit of the *Morinda citrifolia* or noni plant (Tsacas and Bachli, 1981; Louis and David, 1986; Matute and Ayroles, 2014). *D. sechellia* is considered to be a banner species for specialization because its closest relatives are not specialists, and because it relies heavily on *M. citrifolia* at all of its life stages (Louis and David, 1986; R’Kha, Capy, and David 1991; R’Kha *et al*. 1997; Lavista-Llanos *et al*. 2014). Additionally, aside from the ease of cultivating fruit flies in a lab, the ability of *D. sechellia* to hybridize with close relatives facilitated early genetic studies (R’kha *et al*. 1991). Much interest in *D. sechellia* arises from the observation that ripe *M. citrifolia* fruit is highly toxic to other species of fruit flies yet *D. sechellia* is resistant to this toxicity (R’kha *et al*. 1991; Andrade Lopez *et al*. 2017).

The main toxins of *M. citrifolia* fruit are volatile fatty acids, to which *D. sechellia* has evolved both high resistance and preference (Farine *et al*. 1996; Legal *et al*. 1992; Legal *et al*. 1994; Dekker *et al*. 2006; Matute and Ayroles, 2014). A number of studies have centered around the mechanisms of this toxin resistance, most with a focus on the fatty acid volatile octanoic acid (OA) (R’Kha, Capy, and David 1991; Jones, 1998; Lavista-Llanos *et al*. 2014; Lanno and Coolon 2019; Peyser *et al*. 2017; Lanno *et al*. 2017; Lanno *et al*. 2019a). The other primary fatty acid volatile found in noni fruit, hexanoic acid (HA), is also toxic (Farine *et al*. 1996; Peyser *et al*. 2017; Lanno and Coolon 2019) but less so than OA and is responsible for attracting *D. sechellia* to its host (Amalou *et al*. 1998). In that vein, recent work has shown that *D. sechellia* prefers the fruit of *M. citrifolia*, it has adapted to it nutritionally and relies on it for normal reproduction (Lavista-Llanos *et al*. 2014; Watanabe *et al*. 2019).

*D. sechellia* grows and reproduces better on *M. citrofolia* than other food sources in part because of adaptation to a lower carbohydrate to protein ratio (Watanabe *et al*. 2019), but also because of reliance on *M. citrofolia* for L-DOPA, a dopamine precursor (Lavista-Llanos *et al*. 2014). The results of Lavista-Llanos *et al*. explain the observation that maternal environment is more important for larval success in *D. sechellia* than genotype–it is the reliance on an external source of L-DOPA. Surprisingly, they also showed that dopamine confers toxin resistance in other Drosophilids, a result corroborated by Lanno *et al*. (2019a). Additionally, the toxic environment in *M. citrifolia* fruit increases egg production in *D. sechellia* but decreases egg production in other Drosophilae (R’Kha, Capy, and David 1991). A gene expression study using microarrays found that *D. sechellia* fed *M. citrifolia* have increased expression of genes associated with egg production and fatty acid metabolism (Dworkin and Jones 2009). Additionally, *D. sechellia* have low levels of 3,4-dihydroxyphenylalanine (L-DOPA), a dopamine precursor, relative to other *Drosophila* species, likely due a mutation in the *Catsup* gene, which regulates the synthesis of L-DOPA from tyrosine. *M. citrifolia* contains high levels of L-DOPA. When the L-DOPA in *M. citrifolia* is removed, *D. sechellia* produce fewer eggs. Thus, *D. sechellia* supplement their own low levels of L-DOPA with the L-DOPA found in *M. citrifolia* to increase egg production. In addition, this process allows the eggs to survive in the fruit’s toxic environment (Lavista-Llanos *et al*. 2014). Furthermore, *D. melanogaster* and *D. simulans* adult flies fed L-DOPA have increased resistance to OA (Lavista-Llanos *et al*. 2014, Lanno *et al*. 2019a).

In this study we compare transcriptomes of *D. sechellia* fed on a standard diet to a diet supplemented with rotten *M. citrifolia* fruit. The rotten *M. citrifolia* represents a condition in which L-DOPA is preserved and toxicity to other Drosophilids due to OA and HA volatiles is not because microbial action reduces the volatile fatty acids in the fruit (David *et al*. 1989, Lavista-Llanos *et al*. 2014). We also analyze DEGs from each treatment using software that examines shared regulatory motifs among DEGs to make predictions about which transcription factors (TFs) may be regulating the expression of DEGs in response to these different treatments. Therefore, we can make inferences about the role of each compound in altering gene expression in *D. sechellia*, and compare the transcriptomic responses induced by each compound as well as identify regulatory networks that might be common to all *M. citrifolia* related environmental conditions.

## Methods

### RNA-sequencing

Adult female 0-3 day old *D. sechellia* 14021-0428.25 flies were fed control food (Carolina Biological Supply) or control food mixed with 1g rotten *M. citrifolia* fruit pulp for 24 hours. After treatment, flies were snap frozen in liquid nitrogen and stored at −80°C until RNA extraction. Three replicates were analyzed per treatment, with ten flies per replicate, generating 3 control and 3 noni fed samples. RNA was extracted using the Promega SV total RNA extraction system with modified protocol (Promega; Coolon *et al*. 2013). RNA quality was determined using gel electrophoresis and NanoDrop. RNA was sent to the University of Michigan Sequencing Core Facility where mRNA selection was performed from total RNA using poly(A) selection. cDNA libraries were then sequenced using the Illumina Hiseq 4000 platform.

### BIOL310 Genomics Analysis

The genomics analysis of RNA-seq data presented in this manuscript was performed by 2 high school, 20 undergraduate and 3 graduate students as part of a semester-long course at Wesleyan University called Genomics Analysis (BIOL310). This is the fourth such manuscript (see Lanno *et al*. 2017 and Lanno *et al*. 2019a, Drum *et al*. In Review (**BIORXIV/2021/447576)**) generated from this Course-Based Undergraduate Research Experience (CURE) where the aim is to provide early stage undergraduate students an opportunity for hands-on research experience with active participation in the process of scientific discovery. Students in the course learn through engaging with newly generated genomics data and use cutting edge genomics analysis and bioinformatics tools engaging in a discovery-based independent study. Every student in the course contributed to every aspect of the analysis including quality control, bioinformatics, statistical analyses, write-up and interpretation of the findings, providing their own unique perspective of the results and the text written by each and every student was combined into this manuscript with little modification.

After sequencing output files were obtained from the University of Michigan Sequencing Core (Table 1), .fastq files containing raw sequencing reads were uploaded to the Galaxy platform (Afgan *et al*. 2016) and an RNA-seq pipeline analysis was performed (**Figure 1**) as previously described (Lanno *et al*. 2017 and Lanno *et al*. 2019a, Drum et al., In Review (**BIORXIV/2021/447576)**). Briefly, reads were assessed for quality using FASTQC (Andrews 2010) and any overrepresented sequences were analyzed using NCBI Blast (Altschul *et al*. 1990). Bowtie2 was used for mapping reads to the appropriate reference genome for each species with default parameters (Langmead and Salzberg 2012), with the most recent genomes for each species available at the time of analysis acquired from Ensembl (www.ensembl.org, Yates *et al*. 2016) (*D. sechellia:* Drosophila_sechellia.dsec_caf1.dna.toplevel.fa).The Bowtie2 output files were analyzed using Cuffdiff (Trapnell *et al*. 2010), for gene expression quantification and differential gene expression (DEG) analysis using the aforementioned genome file along with the most recent annotated .gff3 file for each genome available at the time of analysis acquired from Ensembl (*D. sechellia*: Drosophila_sechellia.dsec_caf1.42.gff3). In Cuffdiff, geometric normalization and library size correction was performed, along with bias correction using the reference genome, giving an output of DEGs following false discovery rate multiple testing correction (Benjamini & Hochberg 1995, q < 0.05). Data was visualized using R (R Core Team, 2013). The list of DEGs was uploaded to geneontology.org for Gene Ontology term (GO) enrichment analysis (The Gene Ontology Consortium 2000, The Ontology Consortium 2015, geneontology.org). *Drosophila melanogaster* orthologs for each *D. sechellia* gene were downloaded using FlyBase (Thurmond *et al*. 2019). Data processing and visualization was performed in R (R Core Development Team). The *D. melanogaster* orthologs for each DEGs after feeding on noni food were analyzed through the i-*cis*Target analysis software (https://med.kuleuven.be/lcb/i-cisTarget)to identify putative *cis*-regulatory sequences shared among DEGs (Hermann *et al*. 2012, Imrichová *et al*. 2015). The top ten all non-TATA unique sequence elements representing predicted transcription factor binding sites and their downstream targets were then visualized with Cytoscape (https://cytoscape.org/, Shannon *et al*. 2003).). DEGs following *D. sechellia* exposure to OA or L-DOPA were downloaded from the literature (Lanno *et al*. 2017, Lanno *et al*. 2019a). Gene overlap testing was performed using the “GeneOverlap” package in R (Shen 2021) with *D. melanogaster* orthologs.

**Figure 1.**
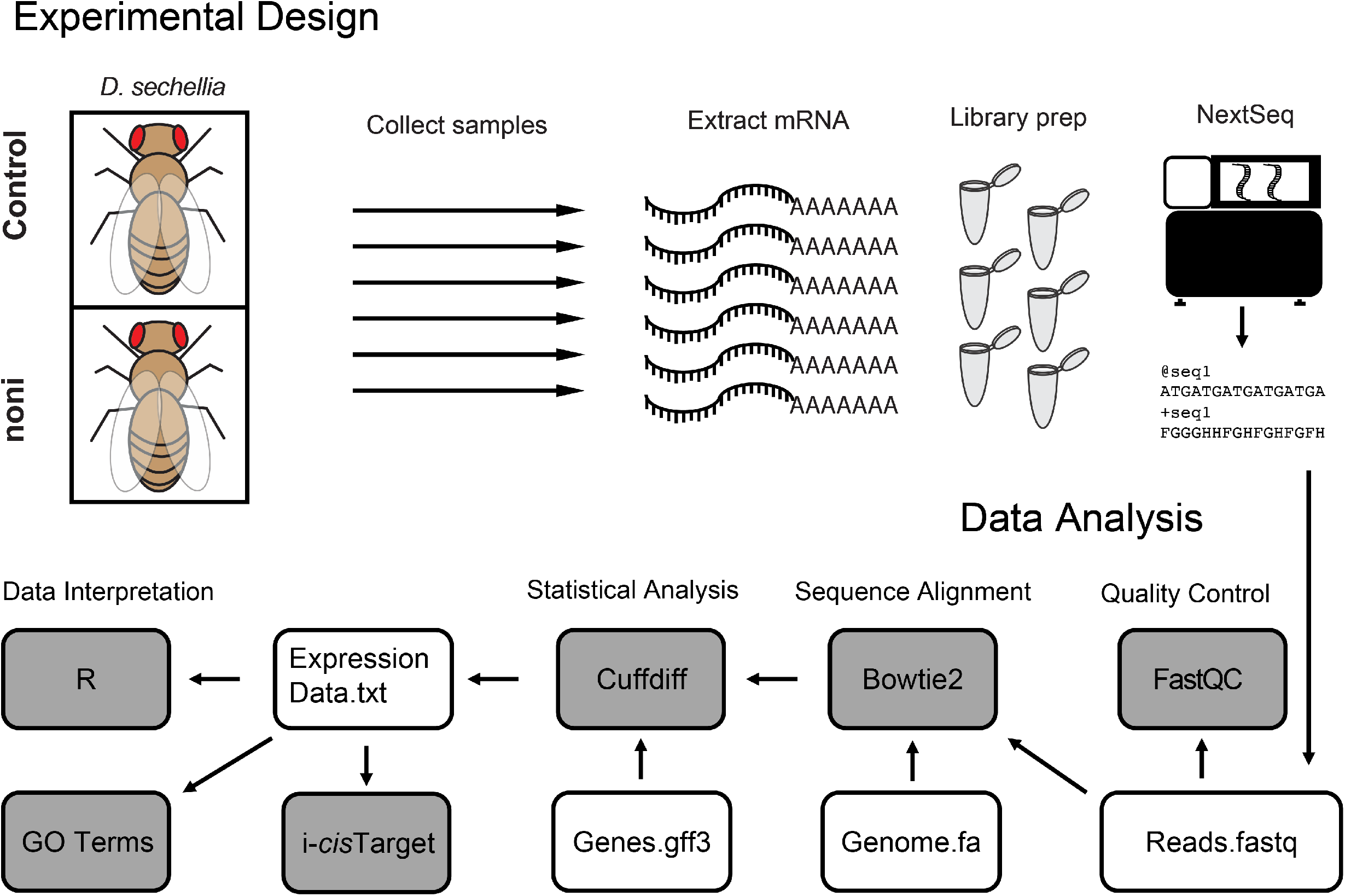
Experimental design. Female *D. sechellia* were exposed to either control food or food supplemented with rotten noni fruit. RNA was extracted, underwent polyA selection, library preparation, and sequencing. Raw sequencing reads were checked for quality using FastQC, and then alligned to the *D. sechellia* reference genome with Bowtie2. Differential expression testing was performed using Cuffdiff, and expression data was analzyed using R, Gene ontology, and i*cis*-Target.

**Table 1:** Samples, mapped reads, and read lengths for each sequencing library.

| Sample | ID | # of Reads | # Mapped Reads | % Mapped | Read length (nt) |
| --- | --- | --- | --- | --- | --- |
| Control 1 | 76332 | 19222060 | 18496450 | 96.23% | 65 |
| Control 2 | 76333 | 20704811 | 19440620 | 93.89% | 65 |
| Control 3 | 76334 | 17696868 | 17123579 | 96.76% | 65 |
| Noni 1 | 76350 | 18937596 | 17939488 | 94.73% | 65 |
| Noni 2 | 76351 | 16870429 | 15801506 | 93.66% | 65 |
| Noni 3 | 76352 | 14595340 | 14181949 | 97.17% | 65 |

### Data accessibility

All RNA-seq data generated in this manuscript have been submitted to the NCBI Gene Expression Omnibus (URL) under accession number (to be available at time of publishing).

## Results

### Identifying genes that are regulated in response to noni fruit diet

To identify the genes that are expressed differently when adult *D. sechellia* flies are fed a diet of control food (instant Drosophila media) compared to flies fed control food supplemented with rotten noni fruit, we used RNA-seq. Statistical analyses of genome wide gene expression in these two diets identified 503 significantly differentially expressed genes (DEGs, **Figure 1**). Of these DEGs, 179 were up-regulated in response to noni fruit and 324 DEGs were down-regulated. Of these 503 DEGs, 421 have annotated *D. melanogaster* orthologs. Of the 82 DEGs without annotated *D. melanogaster* orthologs, 31 DEGs were 5.8S rRNA genes, all of which were downregulated (Table 1, **Figure 2**). Five of the DEGs have uncertain orthologs based on the presence of paralogs in one or more species were removed for further analysis. For the remainder of the analysis, only genes with *D. melanogaster* orthologs were used so annotation for the identified genes could be utilized.

**Figure 2.**
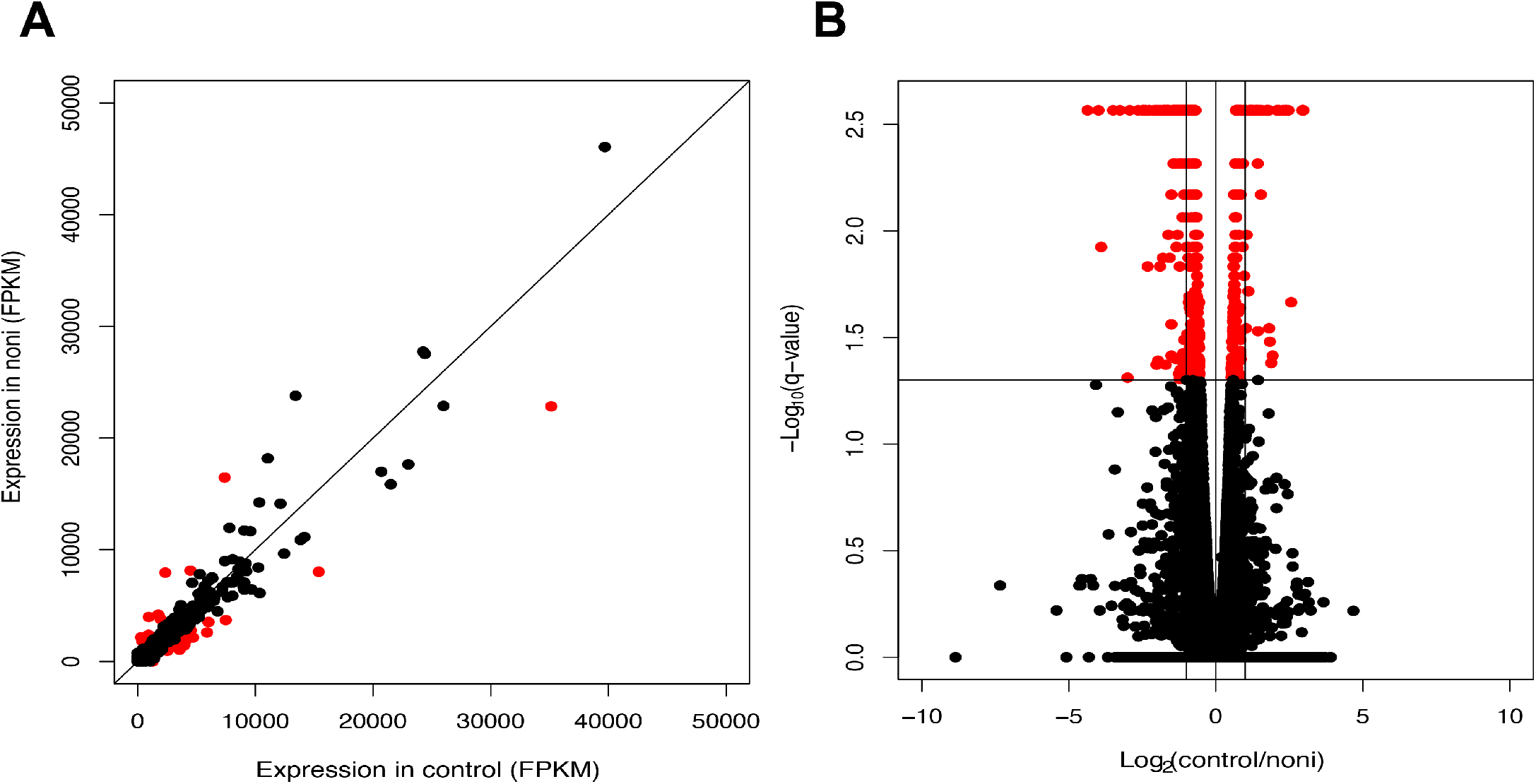
Noni treatment alters genome wide gene expression in adult *D. sechellia*. A. After RNA-seqencing of *D. sechellia* females fed noni fruit, expression of genes (FPKM) in control are shown on the X-axis, with expression of each gene on the Y-axis. Genes that are significantly differentially expressed in noni treatment are shown in red. B. Volcano plot of differentially expressed genes are shown, with Log2(control/noni) on the X-axis, and −Log10(q-value) on the Y-axis. Significantly differentially expressed genes are shown in red.

In order to identify possible functional enrichment among the genes responsive to noni fruit diet, Gene Ontology (GO) term analysis was performed on up- and down-regulated gene sets separately (**Figure 3**). The most significantly enriched biological process GO terms from the up-regulated DEGs included female gamete generation (GO:0007292, =1.32e-08), sexual reproduction (GO:0019953, p=3.06e-07), cell cycle process (GO:0022402, p=8.84-07), chromosome organization (GO: 0051276, p=2.20e-06), chorion-containing eggshell formation (GO:0007304, p=6.26e-06), oogenesis (GO:0048477, p=1.22e-05), cell differentiation (GO:0030154, p=4.27e-05), and epithelial cell development (GO:0002064, p=7.22e-05, **Figure 3A**) consistent with prior genomics and functional studies in *D. sechellia* showing increased egg production in response to feeding on noni (Lavista-Llanos *et al*. 2014; Lanno *et al*. 2019b). The most significantly enriched cellular component GO terms from up-regulated DEGs included egg chorion (GO:0042600, p=3.58e-08), external encapsulating structure (GO:0030312, p=4.43e-08), chromosome (GO:0005694, p=5.04e-06), and non-membrane-bounded organelle (GO:0043228, p=1.15e-05, **Figure 3B**). The most significantly enriched molecular function GO terms from down-regulated genes are alkaline phosphatase activity (GO:0004035, p=2.53e-03), hydrolase activity, hydrolyzing O-glycosyl compounds (GO:0004553, p=5.53e-03), hydrolase activity, acting on glycosyl bonds (GO:0016798, p=8.43e-03) and hydrolase activity (GO:0016787, p=4.66e-02, **Figure 3C**). The most significantly enriched cellular component GO terms from down-regulated DEGs are extracellular region (GO:0005576, p=4.31e-06), cell surface (GO:0009986, p=4.57e-04), plasma membrane (GO:0005920, p=8.77e-03), smooth septate junction (GO:0005920, p=2.83e-02), and nucleus (GO:0005634, p=4.49e-02, **Figure 3D, Tables S7 and S8**).

**Figure 3.**
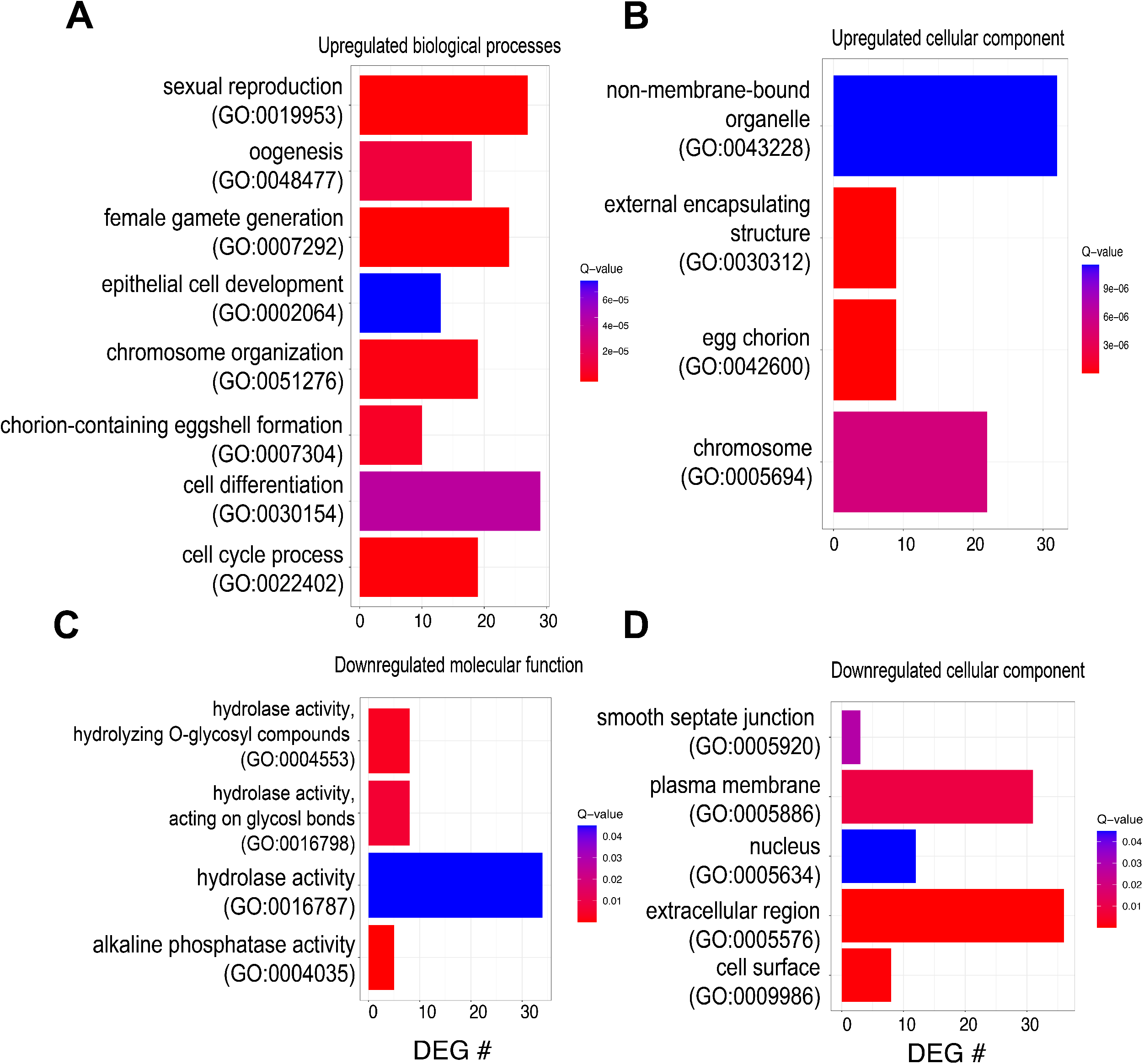
Gene ontology (GO) analysis of differentially expressed genes (DEGs) in response to noni fruit. See table SX for complete lists. (A and B) Upregulated DEGs analyzed for enriched GO processes A: Upregulated DEGs are enriched for several cellular components, including non-membrane-bound organelle, external encapsulating structure, egg chorion, and chromosome. B: Upregulated DEGs are enriched for several biological processes, including sexual reproductive processes, eggshell formation, and cell cycle processes. (C and D) Downregulated DEGs analyzed for enriched GO processes C: Downregulated DEGs and enriched for several cellular components, including smooth septate junction, plasma membrane, nucleus, extracellular region, and cell surface. D: Downregulated DEGs are enriched for several molecular functions, including hydrolase activity and alkaline phosphatase activity.

### Investigating the regulatory network(s) of DEGs responding to noni fruit diet

Identified DEGs were analyzed using i-cisTarget to determine which transcription factors (TFs) may be involved in regulating gene expression upon feeding on rotten noni. Predicted TFs from DEGs responding to noni treatment are *Adf1, GATAd, GATAe, grn, ham, pnr, sd, sim, srp*, and *zld* (**Figure 4**). This analysis predicts all five GATA family TFs to be regulating DEG expression (*GATAd, GATAe, grn, pnr, srp*). GATA factors are important in dietary restriction (Dobson *et al*. 2018) and gut stem cell maintenance (Okumura *et al*. 2015), and may have a role in egg formation in insects (Liu *et al*. 2019) making them excellent candidates for roles in evolved responses to altered diet and downstream effects of diet on egg production. Of the predicted TFs responding to noni treatment, only *sim*, a transcriptional repressor involved in nervous system development is significantly upregulated in *D. sechellia* (Estes, Mosher, and Crews 2001, **Figure S1**.). Of the 31 DEGs predicted to be regulated by *sim* in noni treatment, 23 are downregulated (**Figure 4**).

**Figure 4.**
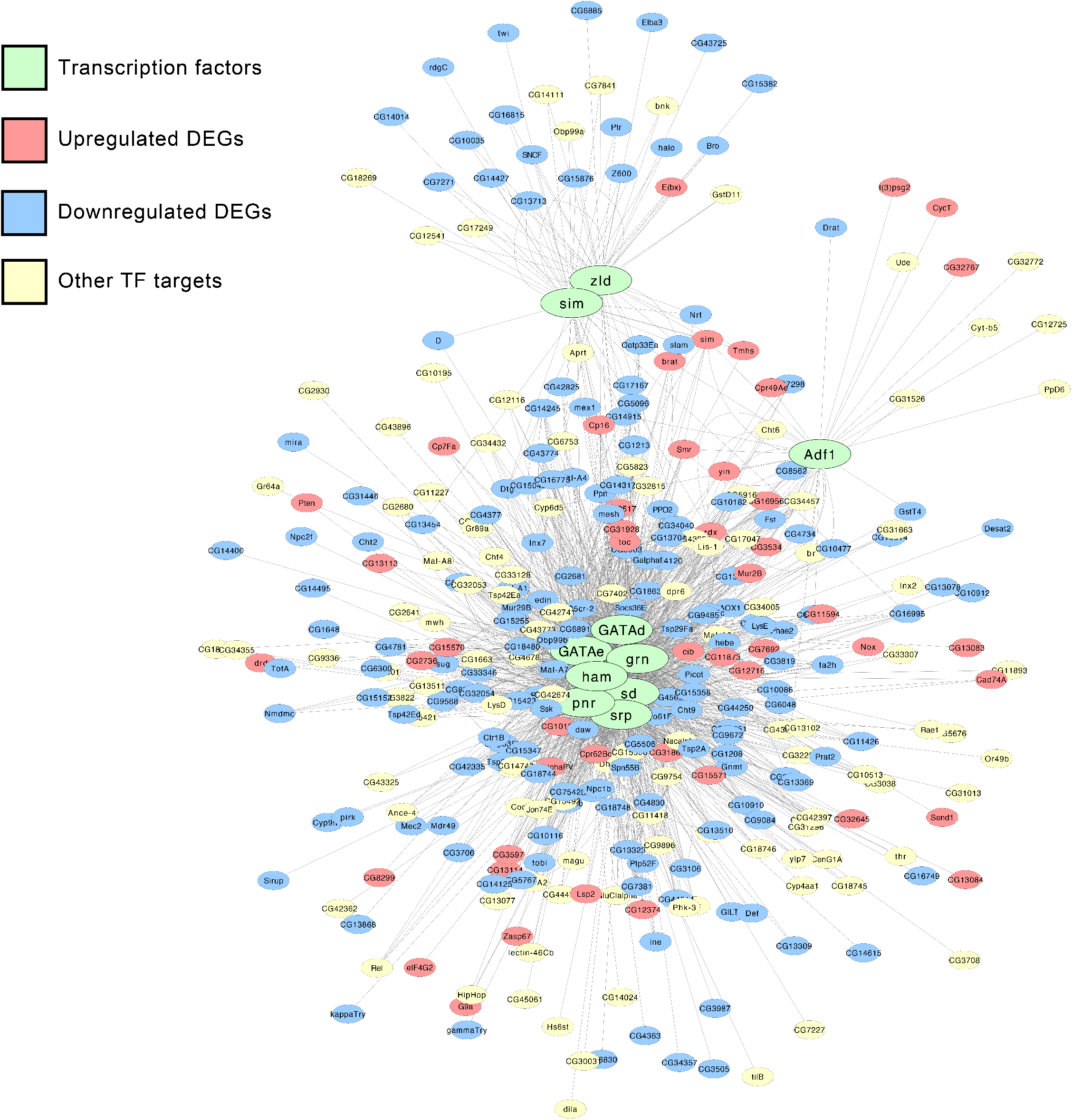
Predicted gene regulatory networks (GRNs) from analysis of significantly differentially expressed genes (DEGs) using i-*cis*Target and visualized using Cytoscape. A. Upregulated DEGs were analyzed and a predicted GRN was assembled, predicting the transcription factors (TFs) *Adf1, EcR, gcm, Hey, Mad, RPII215, Sidpn, sima*, and *sr* (red) to be regulating the expression of upregulated DEGs (purple). Other known targets of these TFs that are not significantly differentially expressed are shown (green). B. Downregulated DEGs were analyzed and a predicted GRN was assembled, predicting the transcription factors *GATAd, GATAe, grn, ham, pnr, sd, sim, srp, zfh1*, and *zld* (red) to be regulating the expression of downregulated DEGs (green). Other known targets of these TFs that are not significantly differentially expressed are shown (orange).

### Comparing DEGs responsive to noni fruit, octanoic acid, hexanoic acid, and L-DOPA

The unique niche that *D. sechellia* utilizes by specializing to feed almost solely on noni fruit, includes multiple highly abundant plant generated chemicals including octanoic acid, hexanoic acid and L-DOPA (Legal *et al*. 1994; Farine *et al*. 1996; Andrade-Lopez *et al*. 2017;). Genome-wide gene expression investigations of responses to each of these individual chemicals were published previously (Lanno *et al*. 2017; Lanno *et al*. 2019a; Drum *et al*. In Review (**BIORXIV/2021/447576)**). Comparing the separate transcriptional responses of *D. sechellia* to the individual chemicals with *D. sechellia* fed noni fruit may help elucidate how they have evolved to specialize on this toxic fruit. Previous studies have investigated the transcriptional response of *D. sechellia* to octanoic acid (OA, Lanno *et al*. 2017), hexanoic acid (HA, Drum *et al*. In Review (**BIORXIV/2021/447576)**), and 3,4-dihydroxyphenylalanine (L-DOPA, Lanno *et al*. 2019a). By analyzing DEGs that do not have annotated *D. melanogaster* orthologs in FlyBase, we found that many are members of different RNA classes. Upon OA, L-DOPA and noni treatment several 5.8SrRNAs, snoRNAs, and 18SrRNAs are all downregulated. In contrast, upon HA treatment several 5.8SrRNAs are upregulated (**Table S3**). Analyzing DEGs in each treatment with annotated *D. melanogaster* orthologs yields 8 DEGs that are significantly differentially expressed in all four treatments and all 8 genes are downregulated (**Figure 5**). The antimicrobial peptides *Defensin, GNBP-like3, edin* are significantly downregulated in all four treatments, as is the transcription factor *Neu2*, as well as other genes: *Sry-alpha, CG14915, CG15876*, and *CG6885*. Gene expression responses in *D. sechellia* exposed to rotten noni or L-DOPA treatment give 173 genes in common. Of these 173 genes, 149 genes are significantly differentially regulated in the same direction in both treatments (Table SX). The transcription factor *sim* is upregulated in both noni and L-DOPA treatments. Interestingly, only 25 of 127 genes differentially expressed in OA treatment (Lanno *et al*. 2017) is specific to only OA treatment and not to other compounds from noni fruit. The two medium chain fatty acids OA and HA share only 2 DEGs (*E(spl)mgamma-HLH and AttA*) between them that are specific to only fatty acid treatment and not L-DOPA or rotten noni (**Tables S1-S4**).

**Figure 5.**
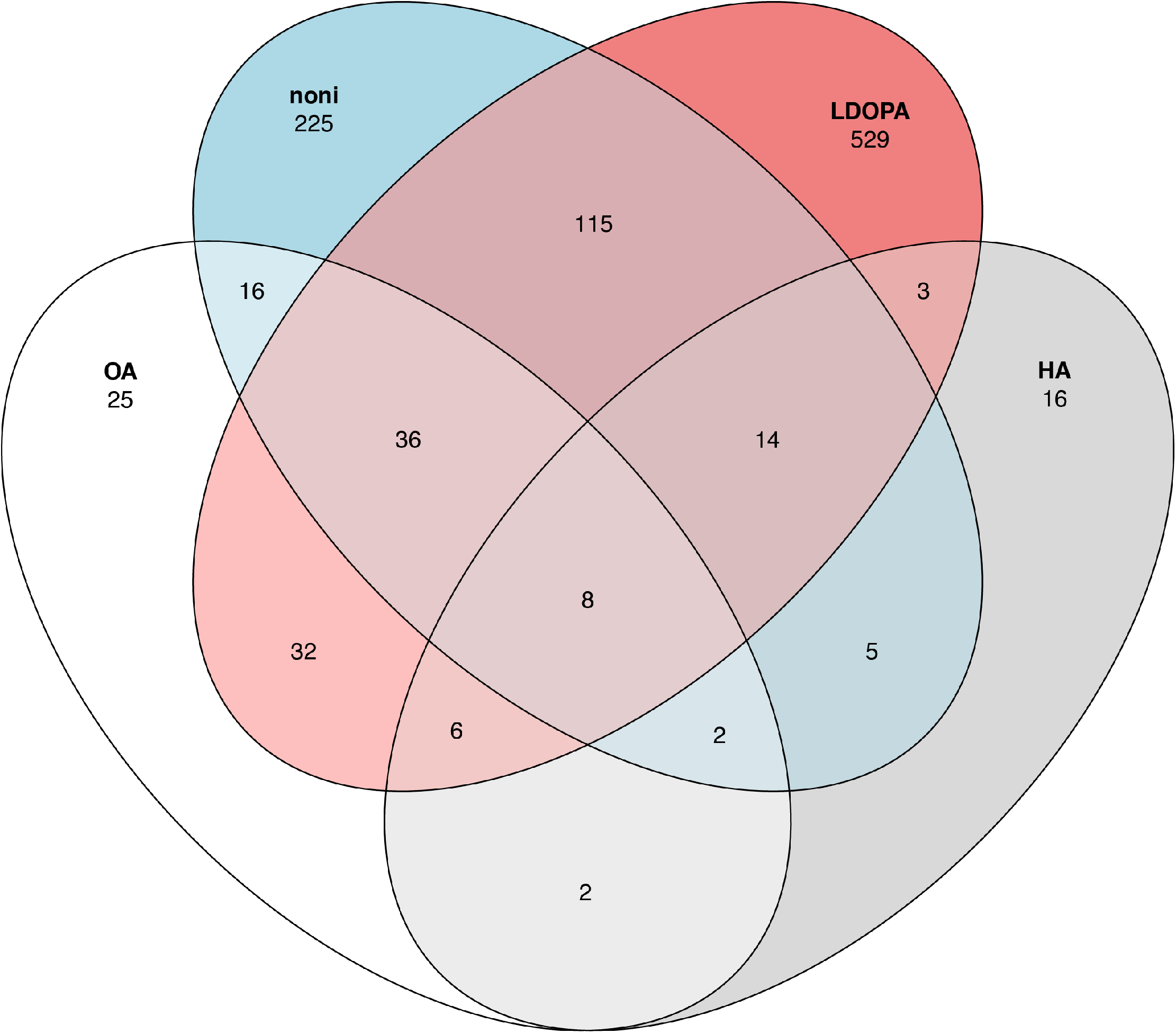
Overlap of significantly differentially expressed genes in response to components of noni fruit. DEGs changing expression in response to OA (white), HA (gray), noni (blue), and L-DOPA (red). Overlaps of shared DEGs are shown, as are DEGs specific to each treatment.

Overlap significance testing was performed by comparing the number of significant DEGs found between each treatment using the “GeneOverlap” package in R (Shen *et al*. 2021). *Drosophila sechellia* has 13,095 genes with annotated *D. melanogaster* orthologs, so this value was used for the genome size measurement. L-DOPA treatment resulted in 743 DEGs, noni treatment gives 421 DEGs, OA treatment resulted in 127 DEGs, and HA treatment resulted in 56 DEGs (**Figure 5**). L-DOPA and noni treatments have 173 overlapping DEGs between these treatments (Fisher’s Exact Test, p=9.8e-108), noni and OA treatments have 62 overlapping DEGs between these treatments (Fisher’s Exact Test, p=5.5e-59), and noni and HA treatments have 29 overlapping DEGs between these treatments (Fisher’s Exact Test, p=6.5e-29). Comparing between previously analyzed treatments, OA and HA treatments have 18 overlapping DEGs (Fisher’s Exact Test, p=2.6e-23), OA and L-DOPA treatments have 82 overlapping DEGs (Fisher’s Exact Test, p=4.3e-71), and HA and L-DOPA treatments have 31 overlapping DEGs (Fisher’s Exact Test, p=1.9e-24). The overlap for every pairwise comparison of DEGs between all four treatments was significant possibly suggesting that there are common regulatory changes that have evolved in *D. sechellia* controlling gene expression responses to multiple aspects of their host food species.

### Comparing predicted TFs among treatments

Significantly differentially expressed genes identified in previous studies that examined OA, L-DOPA, and HA exposure in adult female *D. sechellia* flies (Lanno *et al*. 2017; Lanno *et al*. 2019a; Drum *et al*. In Review (**BIORXIV/2021/447576)**) were used for i-*cis*Target analysis in addition to DEGs we identified here in response to noni to predict transcription factors that control the plasticity of DEGs. For this analysis, all DEGs, both up- and down-regulated are used for all four treatments (OA, L-DOPA, HA and noni). The transcription factor *zelda* (*zld*) was predicted to regulate the expression of genes in all four treatments (**Figure 6**). All 5 GATA family of transcription factors (*grn, pnr, GATAd, GATAe*, and *srp*) were predicted to regulate expression in both noni and L-DOPA treatments, and *srp* was also predicted to regulate expression in HA treatment. *sim* (*single-minded*) was predicted to regulate expression in noni, OA, and HA treatments. The transcription factors *Relish* (*Rel*), *Hsf*, and *Blimp-1* are predicted to regulate expression in both OA and HA treatments. Additionally, *GATAd* expression was significantly increased in L-DOPA treatment, *sim* expression was significantly increased in both L-DOPA and noni treatments, and *dl* expression was significantly increased in L-DOPA treatment (**Tables S1 and S2**). Predicted regulatory networks for each treatment can be found in **Supplemental Figure S2**.)

**Figure 6.**
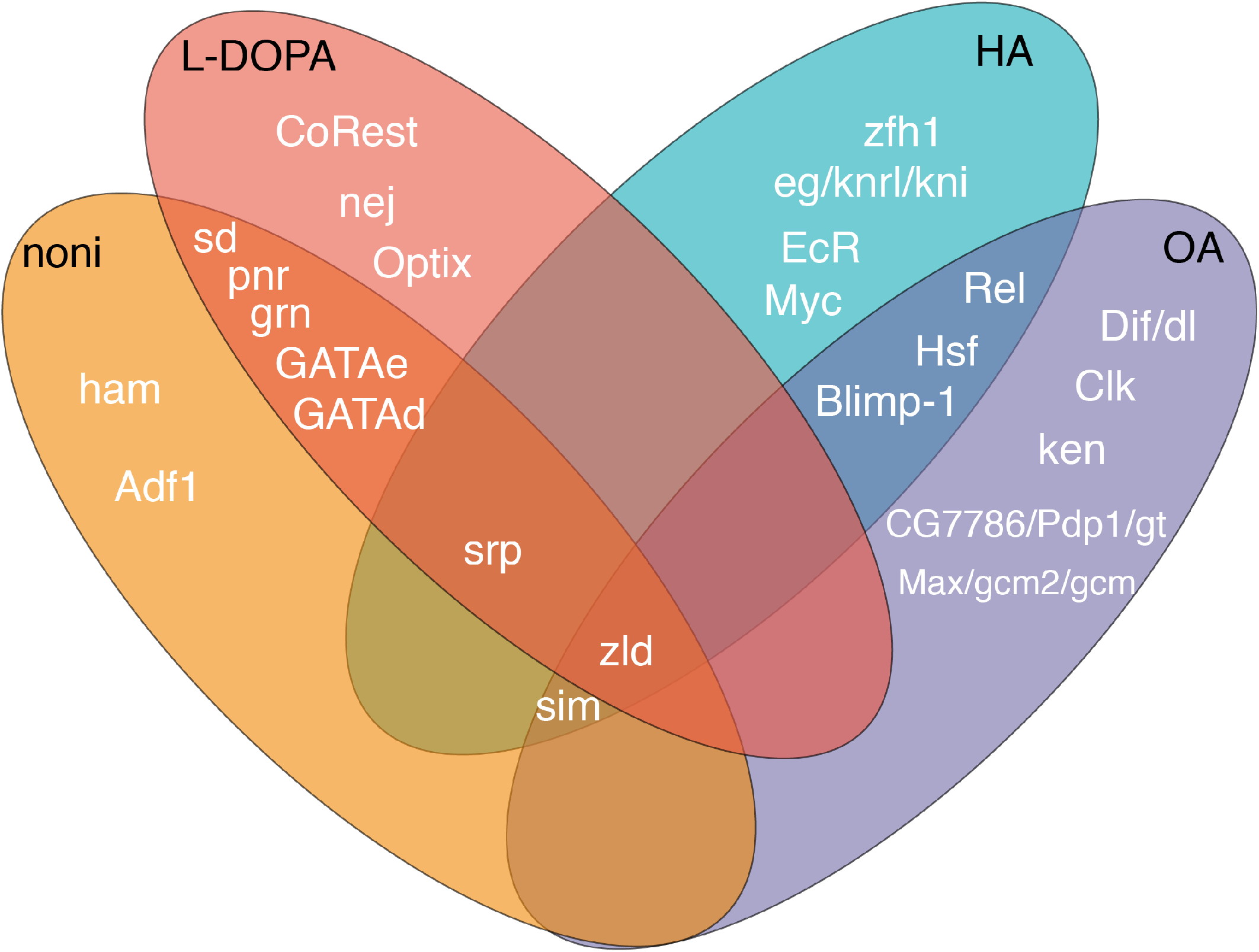
Several transcription factors are predicted to be involved in the response to multiple components of noni fruit. Transcription factors predicted by i-*cis*Target analysis to regulate DEG expression in noni, L-DOPA, HA, and OA treatments in *D. sechellia* are shown. The GATA family of TFs (pnr, grn, GATAe, GATAd, and srp) are predicted to regulate DEG expression in both noni and L-DOPA treatment, with srp also being predicted in HA treatment. Zelda is predicted to regulate expression of DEGs in all four treatments. Single-minded is predicted to regulate DEG expression in noni, L-DOPA, and OA treatment. Rel, Hsf, and Blimp-1 are predicted to regulate DEG expression in both OA and HA treatments. Predicted regulatory networks for each treatment are found in Figure S2.

## Discussion

Examining the whole organism transcriptional response to external influence can help inform the physiology involved in overcoming the stressor (Coolon *et al*. 2009, De Nadal, Ammerer, and Posas 2011. Comparing the physiology that allows *Drosophila sechellia* to feed on toxic *M. citrifolia* fruit to generalist susceptible sister species is an excellent model to study how insects evolve to specialize on toxic resources (Vogel, Musser, and de la Paz Celorio-Mancera 2014). The chemicals found in noni fruit have driven the specialization of *D. sechellia* to its host, as *D. sechellia* has become resistant to the toxic volatile OA, is attracted to the fruit by toxic volatile HA, and utilizes consumed L-DOPA found in noni fruit to facilitate dopamine biosynthesis (Lavista-Llanos *et al*. 2014). Previous studies have examined the whole organism transcriptional response to these separate components of noni fruit that has driven specialization (Lanno *et al*. 2017, Lanno *et al*. 2019a, Drum *et al*. In Review (**BIORXIV/2021/447576)**), and by comparing these transcriptional responses to the transcriptional response to noni fruit alone, we can better understand which responses are specific and which are more general in the evolution of *D. sechellia* noni specialization.

Genes involved in reproductive processes are upregulated in response to noni, L-DOPA, and HA, and genes involved in egg chorion formation are significantly enriched in noni treatment, but significantly downregulated in OA treatment. To better understand how *D. sechellia* has specialized on ripe noni fruit, which contains OA and HA volatiles, understanding the transcriptional response to all components in noni fruit together is necessary. *Drosophila sechellia* selectively oviposits on noni fruit and this increase in proteins involved in egg formation and development are consistent with the evolved reproductive traits for this species when it feeds on noni fruit (R’Kha *et al*. 1997; Lavista-Llanos *et al*. 2014).

The expression of many rRNAs are significantly downregulated in response to both OA and noni treatment, whereas several rRNAs are upregulated in response to HA treatment (Lanno *et al*. 2017; Drum *et al*. In Review (**BIORXIV/2021/447576)**). rRNA synthesis is induced by Ras/Erk signaling in *Drosophila* (Sriskanthandevan-Pirahas, Lee, and Grewal 2018), and Myc and Max are predicted to regulate DEG expression in HA and OA treatments respectively (**Figure 6**). Future work examining how rRNA synthesis and protein translation are involved in the specialization of *D. sechellia* to noni fruit may elucidate a role for these genes in specialization on noni.

Previous studies examining gene expression responses in *D. sechellia* to the volatile fatty acids in noni fruit have focused on examining all of the differentially expressed genes in response to either OA or HA treatment to identify genes involved in evolved toxin resistance (Lanno *et al*. 2017, Lanno *et al*. 2019a, Drum *et al*. In Review (**BIORXIV/2021/447576)**). Interestingly, only 19.7% and 28.9% of DEGs found in these studies are responding to specifically to OA or HA treatment respectively, and only two genes are found that respond to both OA and HA treatment but no other treatments. In order to better understand how insects evolve gene expression responses to plant secondary defenses, it may be helpful to look not only at the response to the toxic chemical, but the wider context of response that more accurately portrays how this interaction would happen in nature where all compounds are experienced simultaneously. The genetic basis of the resistance of *D. sechellia* to OA is polygenic, with the locus conferring the most resistance to OA residing on chromosome 3R (Hungate *et al*. 2013). Previous work has shown that the knockdown of *Osi6, Osi7*, and *Osi8* in adulthood, which reside on this locus, drastically decrease survival to OA (Andrade-Lopez *et al*. 2017). Expression of *Osi6* and several other *Osiris* genes are significantly increased in *D. sechellia* in response to OA (Lanno *et al*. 2017), and *Osi6* is one of the 25 DEGs found only in OA and not in response to any of the other noni components. However, *esterase 6* (*Est-6*) RNAi has been previously shown to alter survival in *D. melanogaster* exposed to OA and the activity of esterase enzymes has previously been shown to be involved in OA resistance (Lanno *et al*. 2017, Lanno *et al*. 2019a, Lanno *et al*. 2019b). Interestingly, *Est-6* expression is significantly enriched in response to L-DOPA and noni treatments, but not upon OA or HA exposure. We propose a mechanism of a gene expression response to the non-toxic chemicals found in the plant that may be priming the gene expression response to help overcome the toxic chemicals found in the plant. As these genes expression response to each of these chemicals evolved together in response to exposure to all components simultaneously, it may be that responses to one chemical confer trait differences important for life in the presence of different components found in noni fruit.

Small changes in transcription factor abundance that alter gene expression in response to external stressor or chemical may be observable by looking at changes in expression of downstream targets of TFs by RNA-seq. Small changes in TF expression are perhaps lost in the error and low sample size of RNA-seq. In order to predict which transcription factors are regulating the transcriptional response to noni in *D. sechellia*, we utilized i-*cis*Target to analyze DEGs to find shared transcription factor binding motifs between DEGs. Expression of each predicted transcription factor was analyzed in response to each treatment, and only a handful are significantly differently expressed between control and treatment measured by RNA-seq (**Tables S1-S4**).

All 5 members of the GATA family of transcription factors are predicted to regulate the expression of DEGs in noni and L-DOPA treatment, along with *srp* being predicted to also regulate DEG expression in HA treatment, but only *GATAd* is significantly upregulated in L-DOPA treatment. U-shaped, the Friend of GATA protein that binds GATA family of transcription factors that has been previously shown to be involved in fatty acid metabolism (Lenz *et al*. 2021) is significantly upregulated in L-DOPA treatment. Expression of GATA factors may be responding to noni and L-DOPA treatment in order to help metabolize volatile fatty acids found in noni fruit that are concurrently experienced by *D. sechellia* in their natural environment when feeding on noni fruit.

*Dorsal* (dl), a TF involved in Toll immune signaling in *Drosophila* (Valanne, Wang, and Rämet 2011), is significantly upregulated only in L-DOPA treatment (**Table S2**). Interestingly, no immune processes are significantly enriched upon L-DOPA exposure in *D. sechellia*, but are significantly downregulated upon OA and HA exposure. Of the genes significantly differentially expressed in all four treatments, many are involved in immune processes. As genes involved in insect immunity are being downregulated in response to OA and HA, future studies examining how the insect microbiome may play a role in the specialization of *D. sechellia* to noni fruit and perhaps resistance to its volatile fatty acids are needed. Gene regulatory response to fatty acids in concert with other components of noni fruit may be involved in regulating the immune system to facilitate the interaction of *D. sechellia* with its toxic host. *Single-minded* (*sim*), a transcription factor that has been previously shown to be a repressor involved in nervous system development (Thomas, Crews, and Goodman 1999; Estes, Mosher, and Crews 2001) is significantly upregulated in both noni and L-DOPA treatments (**Figure S1**). In our network prediction, *sim* is predicted to regulate genes responding to OA, HA, and noni treatments, making it an excellent candidate for a master regulator that evolved to facilitate *D. sechellia* host specialization.

Examining the TFs that are predicted to regulate *Osiris* gene expression may help elucidate how they are regulated in response to OA in *D. sechellia*. From our network analysis of genes responding to OA exposure in *D. sechellia*, the transcription factors *Ken and Barbie* (*Ken*) and *Blimp-1* are predicted to regulate the expression of *Osi6*, one of the *Osiris* genes that was up-regulated upon OA exposure in *D. sechellia* and shown to be involved in resistance to OA toxicity (Lanno *et al*. 2017, Andrade-Lopez *et al*. 2017). More closely examining possible interaction(s) between these transcription factors and *Osiris* genes may shed light on their role in OA resistance.

Separating out genes involved in fruit metabolism compared to genes involved in responding to toxic substances is important for understanding this interaction. Previous work has shown that specialist fruit flies *D. sechellia* and *D. elegans* live significantly longer on protein rich foods than generalist sister species (Watada *et al*. 2020). Noni fruit has a low amount of sugar and is relatively nutrient poor compared to other fruits (Singh *et al*. 2012), so understanding how *D. sechellia* has specialized to eat this fruit and if their transcriptional response plays a role in the metabolism of noni may shed light on how animals alter metabolism to specialize on nutrient poor sources. Examining the potential role of predicted TFs and other DEGs may help us understand the transcriptional response of *D. sechellia* to noni fruit and shed light on the genetic basis of *D. sechellia* evolved specialization on its toxic host plant. Using network prediction tools to understand the regulatory environment of gene expression may elucidate how gene regulation is being altered that would be missed in the analysis of differentially expressed genes alone, especially when there are hundreds of DEGs to analyze. Using a combination of transcriptome sequencing with methods to predict which transcription factors are responsible for gene expression responses due to external compounds may elucidate how insects are able to adapt to harsh environments and evolve to specialize on new and frequently toxic hosts plant species.

**Table 2:** DEGs in *D. sechellia* in response to noni treatment.

| # DEGs | # Upregulated | #Downregulated | # of *D. melanogaster* orthologs |
| --- | --- | --- | --- |
| 503 | 179 | 324 | 426 |

---

Figure S1, Change in expression of predicted TFs in response to noni treatment. A. Transcription factors predicted by i-*cis*Target to regulate the expression of upregulated DEGs in noni treatment and the mean of the log_2_ Fold Change is shown. None of these transcription factors are significantly differentially expressed in response to noni treatment. B. Transcription factors predicted by i-*cis*Target to regulate the expression of downregulated DEGs in noni treatment and the mean of the log_2_ Fold Change is shown. *Sim* is the only of these transcription factors found to be significantly differentially expressed in response to noni treatment.

Figure S2. Predicted gene regulatory networks for each treatment are shown. Predicted transcription factors are shown in green, upregulated DEGs are shown in red, downregulated DEGs are shown in blue, and other targets of these TFs are shown in yellow.

## Acknowledgements

Research reported in this publication was supported by Wesleyan University (Startup to JDC, Department of Biology funds to JDC, College of the Environment funds to JDC), and the National Institute Of General Medical Sciences of the National Institutes of Health under Award Number R15GM135901 (award to JDC). The content is solely the responsibility of the authors and does not necessarily represent the official views of the National Institutes of Health.

